# Large-Scale Information Retrieval and Correction of Noisy Pharmacogenomic Datasets through Residual Thresholded Deep Matrix Factorization

**DOI:** 10.1101/2023.12.07.570723

**Authors:** Zhiyue Tom Hu, Yaodong Yu, Ruoqiao Chen, Shan-Ju Yeh, Bin Chen, Haiyan Huang

## Abstract

Pharmacogenomics studies are attracting an increasing amount of interest from researchers in precision medicine. The advances in high-throughput experiments and multiplexed approaches allow the large-scale quantification of drug sensitivities in molecularly characterized cancer cell lines (CCLs), resulting in a number of open drug sensitivity datasets for drug biomarker discovery. However, a significant inconsistency in drug sensitivity values among these datasets has been noted. Such inconsistency indicates the presence of substantial noise, subsequently hindering downstream analyses. To address the noise in drug sensitivity data, we introduce a robust and scalable deep learning framework, Residual Thresholded Deep Matrix Factorization (RT-DMF). This method takes a single drug sensitivity data matrix as its sole input and outputs a corrected and imputed matrix. Deep Matrix Factorization (DMF) excels at uncovering subtle patterns, due to its minimal reliance on data structure assumptions. This attribute significantly boosts DMF’s ability to identify complex hidden patterns among nuisance effects in the data, thereby facilitating the detection of signals that are therapeutically relevant. Furthermore, RT-DMF incorporates an iterative residual thresholding (RT) procedure, which plays a crucial role in retaining signals more likely to hold therapeutic importance. Validation using simulated datasets and real pharmacogenomics datasets demonstrates the effectiveness of our approach in correcting noise and imputing missing data in drug sensitivity datasets (open source package available at https://github.com/tomwhoooo/rtdmf).

## 1. INTRODUCTION

One goal of precision medicine is to choose the best therapy for individual cancer patients on the basis of individual molecular markers identified in clinical studies (as in Collins and Varmus (2015); Lowy and Collins (2016); Chen and Butte (2016)). At present, only some cancer drugs have approved biomarkers, and the process of identifying and validating a biomarker for a single drug in clinical trials takes many years (Yothers *and others* (2013); de Gramont *and others* (2015)). Emerging pharmacogenomics studies, in which drugs are tested against panels of molecularly characterized cancer cell lines (CCL’s), have enabled the identification of several types of molecular biomarkers on a large scale by correlating drug sensitivity with the molecular profiles of pre-treatment cancer cell lines (Garnett and et al. (2012); Barretina *and others* (2012); Basu *and others* (2013); Iorio *and others* (2016); Niepel *and others* (2013)). These biomarkers have the potential of identifying cell lines or even patients that will respond to a particular drug.

The Broad Institute’s Cancer Cell Line Encyclopedia (CCLE) project released the CCLE dataset in 2012. This dataset comprised sensitivity data for 1046 cell lines across 24 compounds (Barretina *and others* (2012)). Around the same period, the Sanger Institute’s Genomics of Drug Sensitivity in Cancer Genome Project introduced the GDSC dataset. This set covered sensitivity data from nearly 700 cell lines and 138 compounds (Yang *and others* (2013)). Since the release of both CCLE and GDSC, several similar pharmacogenomic studies have been published. A subsequent dataset from the Broad Institute’s Cancer Therapeutics Response Portal project, known as CTRPv2, encompassed 860 cell lines and 481 compounds (Rees *and others* (2016); Seashore-Ludlow *and others* (2015); Basu *and others* (2013)). The updated version of the Genomics of Drug Sensitivity in Cancer dataset, termed GDSC1000, includes 1001 cell lines and 251 compounds. Additionally, there have been pharmacogenomics studies that are specific to particular types of cancer, such as acute myeloid leukemia (Marcotte *and others* (2016); Daemen *and others* (2013); Lee *and others* (2018)).

The datasets mentioned above have made important contributions to pharmacogenomics research. Each drug sensitivity dataset can be represented as a single data matrix, where each row denotes a drug, each column symbolizes a cancer cell line, and the values — often interpreted as drug responses — are summarized from dose-response curves. Commonly employed summarization metrics of the dose-response curves include IC50 (the concentration at which the drug inhibits 50% of maximum cell growth) and AUC (area under the dose-response curve). However, some of these matrices may contain a lot of missing data or incorrect zeros, making them difficult to use in subsequence analysis. Furthermore, re-evaluations of previously published pharmacogenomics data have exposed inconsistencies in drug response data across different studies. A comparative study by Haibe-Kains *and others* (2013) between two datasets, CGP and CCLE, focusing on 15 shared drugs tested on 471 common cell lines, found that the vast majority of drugs displayed poor concordance. This inconsistency raises concerns regarding their applicability in biomarker discovery and brings into question the reliability of such data for drug discovery, as noted in Haibe-Kains *and others* (2013) and Safikhani *and others* (2016).

Numerous attempts have been made to address this inconsistency problem. Most of the proposed ideas focused on forming better summarization metrics and/or standardizing experiments and data processing pipelines (e.g., Mpindi *and others* (2016); Bouhaddou *and others* (2016); Hafner *and others* (2016)). Although these methods have improved the cross-study consistency of drug response measures to some extent, there is still a vast inconsistency of drug response signals across the studies. The study by Hu *and others* (2019) proposed a novel semi-supervised computational method named AICM, designed to correct sensitivity data matrices. This method was inspired by an intriguing observation: when comparing the same cancer cell lines across different datasets, the results are usually consistent, with high correlations. However, this consistency is not observed when examining the effects of the same drug across different datasets, where correlations are low. These findings suggest that while there is some level of agreement between various studies, there’s also considerable noise or error in the data. AICM leverages the consistent correlations in cancer cell lines to correct and impute missing data across different datasets. A limitation of this approach is that it requires datasets with a substantial overlap in both drugs and cancer cell lines to be effective. As newer studies increasingly focus on specific types of cancer or particular drug libraries, this limitation becomes more significant. Therefore, rather than relying on shared information from overlapping sections of datasets, it’s essential to develop a method that individually corrects each dataset by thoroughly examining its underlying structure. This approach should surpass traditional methods by capturing subtle and intricate patterns and signals. Additionally, it must accommodate large amounts of missing data and be scalable to larger datasets.

In this paper, we present Residual Thresholded Deep Matrix Factorization (RT-DMF), a novel computational framework that operates in a fully unsupervised manner and aims to reveal hidden signals within individual datasets. RT-DMF is developed with the understanding that therapeutically significant interactions between drug and cancer cell lines (CCLs) are sparse in drug sensitivity data matrices, leading to the assumption that most patterns identified by matrix factorization methods are likely to represent nuisance drug-CCL relationships. This framework takes a single drug sensitivity data matrix as its sole input and outputs a corrected and imputed matrix. RT-DMF has two key features: i) It advances traditional matrix factorization methods by employing a linear neural network. This eliminates the need to assume that the data matrix for nuisance effects is the inner product of two low-rank matrices, thereby allowing for a more effective representation of complex nuisance structures or patterns in the data. This, in turn, aids in the detection of signals that are more therapeutically relevant. ii) It incorporates a residual filtering process to recover valuable data that might otherwise be categorized as outliers. Through extensive experiments, we have demonstrated that RT-DMF not only denoises individual datasets but also enhances consistency across multiple datasets. Furthermore, we show that the method’s capability to impute missing values has significant benefits for various downstream analyses. Based on these results, we believe that RT-DMF holds the potential to become a standard processing pipeline for future datasets.

## 2. METHOD

### 2.1 Method overview

Modeling the collaborative relationships between drugs and Cancer Cell Lines (CCLs) is structurally and numerically complex, regardless of whether these relationships are nuisance or therapeutically interesting. In ideal situations with discernible patterns in datasets, we may observe randomly distributed “blocks.” These blocks suggest scenarios where a group of drugs similarly affects a set of CCLs. The size of these groups can vary; for instance, non-selective drugs active across many cell lines form larger blocks, possibly indicating nuisance drug-cancer relationships. On the other hand, specialized drugs might be effective on a specific small subset of CCLs, resulting in smaller blocks, potentially signifying therapeutically relevant relationships.

Moreover, some drugs may have multiple modes of action, targeting different CCL groups, while certain CCLs could be influenced by diverse drug groups acting on distinct pathways. This implies that some drugs or CCLs might belong to multiple groups or blocks. “Stripes” may also appear, for example, from cytotoxic drugs that show efficacy across most CCLs. Additionally, hidden non-block patterns and various types of noise likely exist. For instance, a group of drugs targeting a common mutation in CCLs may show inconsistent efficacy due to each drug’s unique and intricate mechanism of action.

Explicit models face challenges in effectively capturing the above patterns and differentiating between nuisance and therapeutically relevant drug-CCL relationships, particularly in the presence of data noise. Therefore, a model with fewer structural assumptions but greater capability to uncover hidden nuisance patterns, while remaining robust against noise, is considered to have more promise for this task.

Considering that therapeutically relevant drug-CCL relationships are sparse in a drug sensitivity data matrix, it is reasonable to assume that patterns identified through matrix factorization methods would mostly represent nuisance drug-CCL relationships. In this study, we introduce a new method based on deep matrix factorization (DMF) to isolate these nuisance patterns within drug sensitivity data. We compared DMF with classical low-rank matrix factorization methods and found that the DMF-based approach has advantages in extracting these nuisance patterns. This extraction can, in turn, facilitate the identification of signals that are more therapeutically relevant.

Classical low-rank matrix factorization aims to approximate a data matrix *𝒟 ∈* ℝ^*n×p*^ as *𝒟 ≈ M*_1_*M*_2_, under the assumption that *M*_1_, *M*_2_ are two low-rank matrices. While this approach can be useful and provide interpretable patterns for straightforward interactions, it falls short when faced with more complex data structures, as described earlier regarding possible groups and blocks.

The primary motivation for DMF lies in its ability to combine (i) the interpretability and robustness found in classical matrix factorizations, with (ii) the capacity to extract patterns of varied signal levels and intricate structures as enabled by multi-layer architectures. Single-layer matrix approximations struggle to capture patterns with widely varying signal levels in complex datasets. DMF decomposes a data matrix *𝒟 ∈* ℝ^*n×p*^ as *𝒟 ≈ M*_1_*M*_2_ … *M*_*L*_, where *L* represents the number of layers. The approximation achieved by DMF can be understood as a series of successive factorizations of *𝒟*:

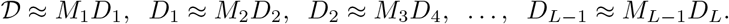

Here, each matrix *M*_*l*_(*l* = 1, …, *L*) can be interpreted as the feature matrix of layer *l*. Consequently, the matrix *D*_*l*_(1 ⩽ *l* ⩽ *L*) is successively factored so that key linear combinations of the principal features of the *l*-th layer appear in the subsequent *M*_*l*+1_. This enables a wealth of interpretations concerning the semantics concealed within the data.

Another key feature of the DMF approximation is its optimization approach. By initializing the matrices *M*_*l*_ (where *l* = 1, …, *L*) as high-dimensional matrices, DMF minimizes the assumptions it makes about the underlying structure. This, however, introduces the challenge of navigating an unidentifiable search space. Despite this obstacle, the use of gradient descent methods effectively regularizes the solution path, enabling DMF to autonomously arrive at a lowrank version of *M*_*l*_’s. This phenomenon of implicit regularization – where the model optimizes itself towards a simpler, more generalizable solution – is well-studied, both theoretically and experimentally, in various research papers (e.g., Neyshabur (2017); Arora *and others* (2019)).

In contrast to classical matrix factorization methods, which operate under stronger structural assumptions and have fewer parameters, thereby increasing the likelihood of achieving global optimization, DMF offers potentially greater fidelity to the underlying data. This is because it operates in a more relaxed, larger search space. Although global optimization is unlikely achievable, the method’s greater flexibility increases the odds of finding solutions that are closer to the true underlying structure.

After applying DMF to a drug sensitivity data matrix, as we argued earlier, we assume that the DMF approximation captures structured nuisance signals. The residuals are likely composed of remaining random or unstructured nuisance noise, as well as “true” or therapeutically relevant signals. In different applications, significant deviations from DMF approximations might be dismissed as outliers or noise. However, in the context of drug sensitivity data, these substantial deviations could indicate important signals. For instance, a small number of tested CCLs might have a unique mutation that is specifically targeted by a drug, leading to “spikes” in the matrix. To preserve this crucial information, we incorporate a residual thresholding (RT) step. This process starts with identifying high residuals, marked by deviations exceeding a pre-determined threshold. We then examine if the CCLs associated with these high residuals under the same drug treatment are over-represented in known CCL groups, such as those categorized by tissue type, cancer type, or cancer subtype. We base this approach on the hypothesis that high residuals enriched within specific tissue or disease groups are more likely to hold therapeutic significance.

In summary, RT-DMF decomposes the signals in a drug sensitivity data matrix by: I) using DMF to extract and remove structured nuisance signals; II) applying thresholding to the residuals to separate out potentially therapeutically significant signals; III) within the thresholded residuals that show substantial deviations from DMF approximations, pinpointing those that are prevalent in known tissue or disease categories. Signals identified in Step III are regarded as having therapeutic interest. The final output is then the DMF approximation matrix with the high residuals identified in Step III added back to the corresponding entries.

For the step III, in situations where reliable CCL groupings are unavailable, we suggest a more straightforward method to retain these notably deviating values: preserving values that show abrupt changes in a single training iteration. We provide more details on this methodology in the subsequent algorithm section.

### 2.2 Algorithm

Building on the ideas outlined in the previous section, the RT-DMF algorithm is presented below. More details regarding the implementation of RT-DMF can be found in the github repository.

#### Algorithm 1

Deep Matrix Factorization with Residual Filtering

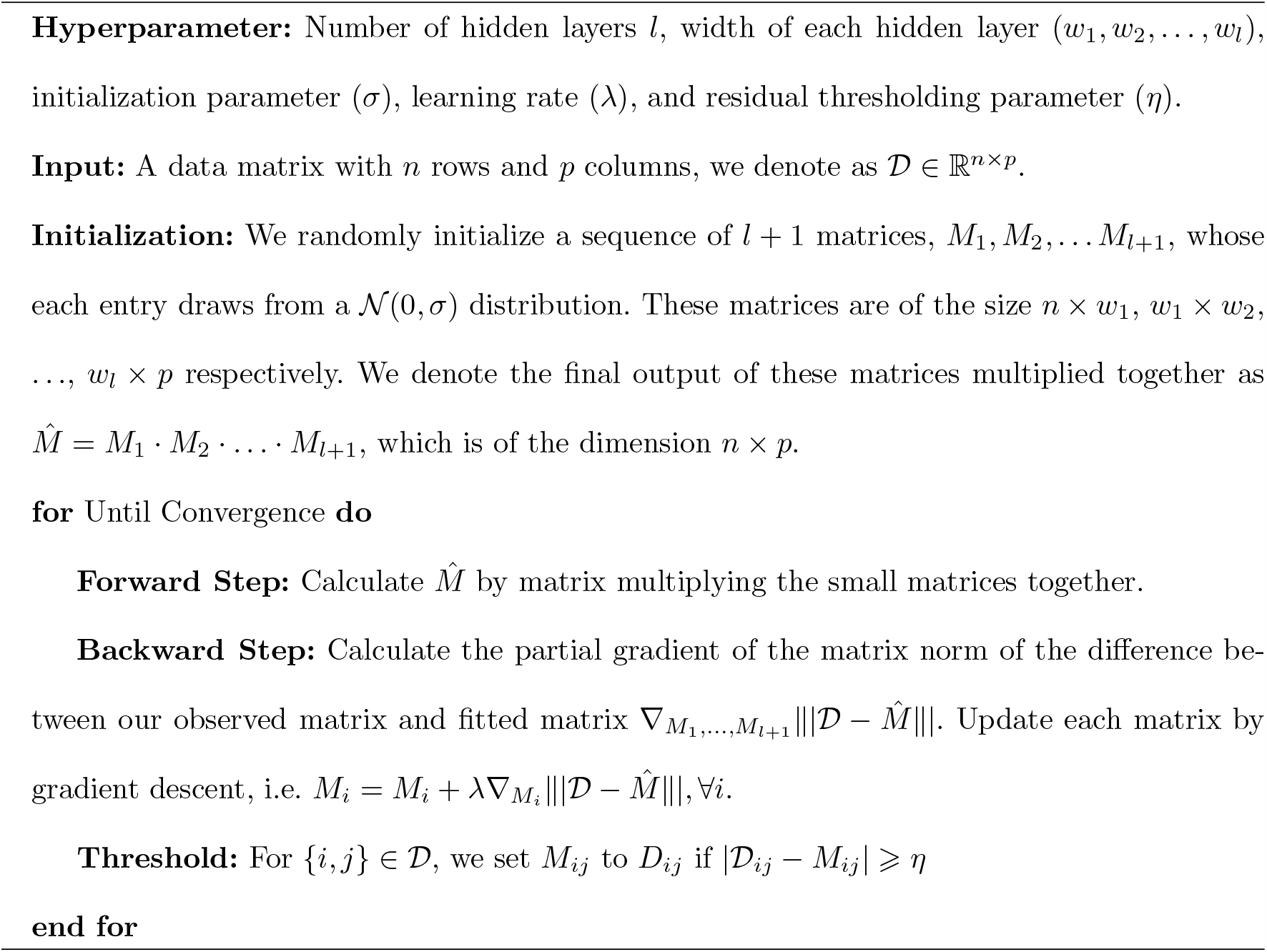

## 3. SIMULATION RESULTS

### 3.1 Overview

Unless otherwise specified, we denote a particular drug sensitivity matrix as *𝒟 ∈* ℝ^*n×p*^, meaning that this dataset has *n* drugs tested on *p* CCLs. We denote the row index of *𝒟* as *r*_1_, *r*_2_, …, *r*_*n*_ and column indices as *c*_1_, *c*_2_, … *c*_*p*_. *𝒟* [*i, j*] denotes the single entry on the *i*-th row and *j*-th column of the matrix *𝒟*, which is a summarized value of dose-response of drug *i* on CCL *j*.

Given that our ultimate goal is to uncover signals in an unsupervised manner, we attempt to generate synthetic datasets that incorporate the complexity of drug-CCL interactions as we previously discussed to verify the effectiveness of the methods. We evaluate the performance of RT-DMF on these synthetic datasets by comparing it with the most widely-used matrix factorization methods: Convex Matrix Completion in Candès and Recht (2009) and Robust Principal Component Analysis (RPCA) in Candès *and others* (2011). Note that the same residual thresholding techniques are used in these benchmark methods for fair comparison.

We generate our synthetic data using the following scheme:

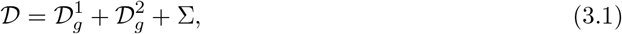

where 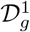 and 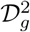 denote different kinds of “structural information”. 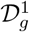 denotes more general interaction information, such as the effect of general type/group of drug(s) on a relatively large group of CCLs, thereby representing nuisance drug-CCL relationships. More specifically, we model 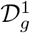 as a low-rank matrix that can be recovered as the product of multiple matrices. As we discussed earlier, modeling the nuisance interactions between drugs and CCL as the simple inner product of two matrices would be an oversimplification because such interactions are complex. Therefore, we parameterize it as a multiplication of multiple wide matrices to mimic a dataset with patterns of varied signal levels and complicated structures. Nevertheless, we also provide a case where 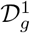 is a simple multiplication of two low-rank matrices, which is consistent with the assumptions of other benchmark methods. We model 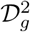as comprising combinations of “extreme values.” These effects may result from a drug’s strong toxicity on the majority of CCLs, which can create a row stripe in the matrix. Additionally, they can arise from signals corresponding to more specific drug-CCL interactions, which are distinct from the nuisance effects in 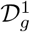 and may appear as tiny individual blocks or overlapping small blocks in the matrix. In other words, 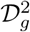 can consist of high residuals that are enriched in specific tissue or disease groups of CCLs (“extreme values”), indicative of signals with greater therapeutic interest. We will introduce various models of 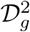 in the next subsection.

In summary, as previously discussed, we anticipate that 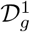 will represent nuisance interactions between drugs and CCLs. In contrast, 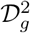 contains information about more specific or extreme drug-CCL interactions. Specific drug-CCL interactions are likely to be therapeutically relevant. It’s important to note that accurately recovering 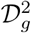 depends largely on the successful recovery of 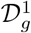.

Σ represents the noise introduced either by experimental protocols or other lab environmental factors. In the RT-DMF framework, DMF is utilized to effectively recover 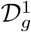. The RT procedure is then employed to recover 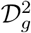 from the residuals. The remaining residuals account for the noise as described in Σ. Generally, separating 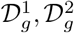 and Σ is a challenging task. As we will demonstrate later, careful tuning of parameters can provide a viable approach to achieve this separation.

### 3.2 Generation of the synthetic datasets

We outline the generation models for 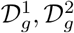 and Σ below, which have dimensions of 800 *×* 400 (i.e., 800 drugs and 400 CCLs). Detailed information about the generation process can be found in the Appendix.

- **Generation of** Σ. Each component in Σ is sampled from a normal distribution *N* (0, *σ*^2^).
- **Generation of** 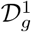. We generate 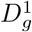 using two models: the first is a product of two nonnegative low-rank matrices, while the second is a product of multiple, such as four, low-rank matrices.
- **Generation of** 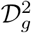. We consider three different block structures to generate 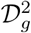: (i) **Simple two-block**. This models an ideal situation where a small group of drugs appears effective for a small group of CCLs. We generated two blocks, each with dimensions of 20 *×* 20, with non-overlapping drugs or CCLs. A zoomed-in visualization of this block structure within a submatrix of 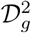 is shown in Figure 1d. Prior to applying RT-DMF, we shuffled the indices to ensure that the blocks are not immediately apparent in the data. (ii) **Mixed overlapping blocks**. This models more complex scenarios in which CCLs may respond to different drug mechanisms, resulting in blocks with overlapping CCLs as depicted in Figure 2d. In addition, we consider the case where there are possible big categories of cancer: that certain CCLs might respond to almost all drugs, resulting in vertical stripes in the data representation (see column 15 in the figure). (iii) **Real data residual**. This model generates a synthetic data matrix by utilizing clustered residuals from the FIMM data (Mpindi *and others* (2016)), combined with the naive low-rank approximation of the FIMM data matrix. This provides a simplified representation of real data, but with the structures of both nuisance and potentially therapeutic signals known. A visualization of a sub-matrix of 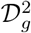 can be seen in Figure 3d.

**Fig. 1:**
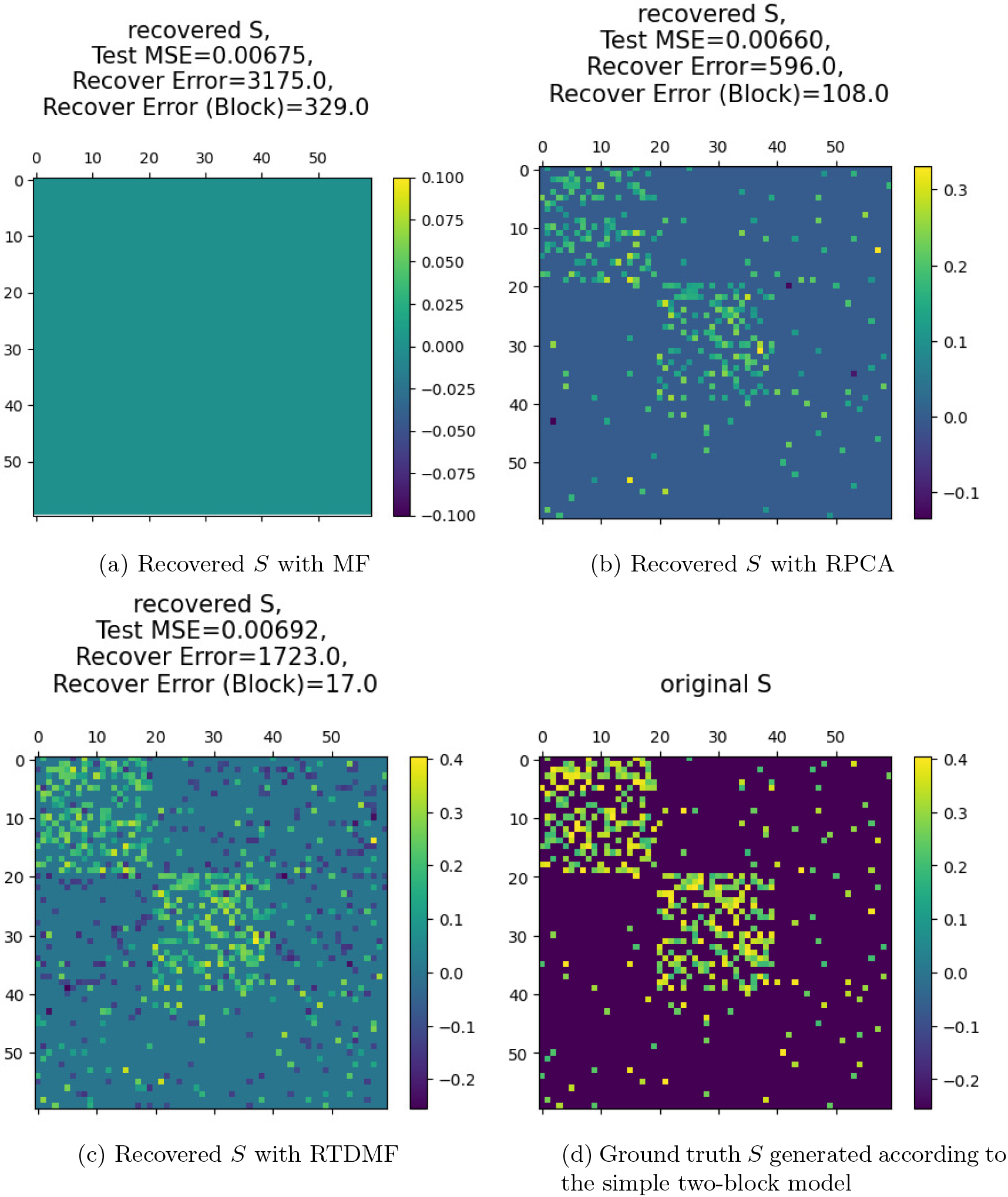
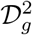 model 1: simple two-block model

**Fig. 2:**
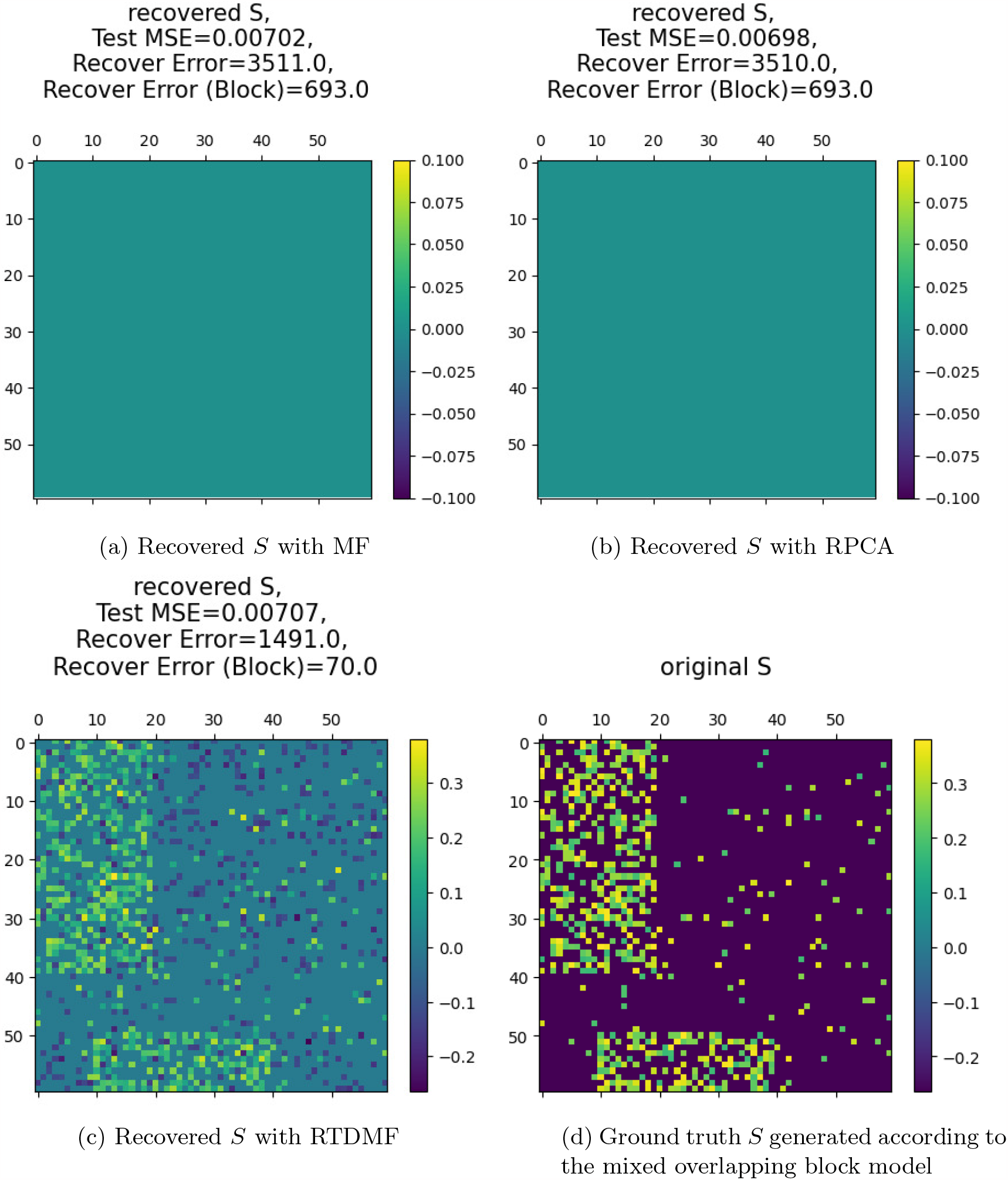
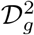 model 2: mixed overlapping block model

**Fig. 3:**
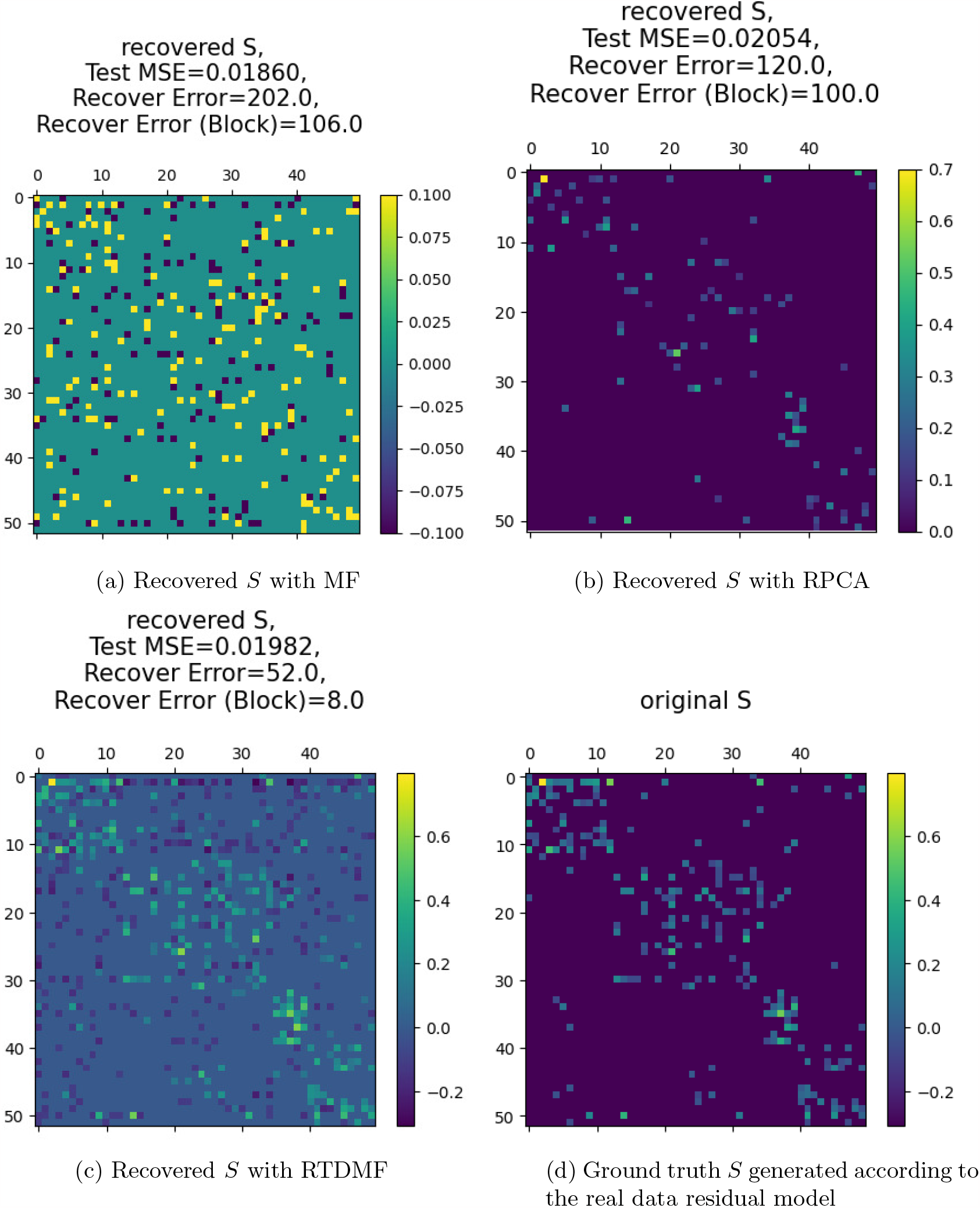
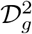 model 3: residual fetched from co-clustered FIMM residual

### 3.3 Comparison methods and results

We compare RT-DMF with benchmark methods including the simple convex matrix factorization (MF) proposed by Candès and Recht (2009) and the Robust Principal Component Analysis (RPCA) proposed by Candès and others (2011). A more comprehensive description of these two methods can be found in the appendix.

For simple MF, the crucial tuning parameter is *ϵ*, which controls the similarity between the fitted and original matrices and determines the rank of the fitted matrix. In RPCA, the key tuning parameter is *τ* ; as *τ* increases, the fitted matrix becomes sparser. For RT-DMF, we adjusted the number of epochs (with a fixed learning rate), balancing the fit to the original matrix against retaining structural low-rank information. As the number of epochs increases, the matrix recovered through RT-DMF has a higher rank to better fit the original matrix. All these parameters represent a trade-off between training error and the assumed structural sparsity.

For all three methods, we tune the parameters based on the validation mean square error (v-MSE). Specifically, we use only 70% of the data for training. The remaining 30% of the data in the matrix is not utilized during the optimization process. Of this, 15% is used as validation entries for parameter selection, and the other 15% serves as test entries for evaluating the model’s performance using the test MSE. We select the parameter that yields the best v-MSE for testing purposes.

We assess the performance of the methods using three different metrics: test mean square error (MSE), recovery error (RE), and in-block recovery error (IBRE). Test MSE refers to the MSE calculated on the test entries. RE quantifies the overall difference between the recovered matrix and the original matrix. IBRE measures the difference specifically for entries that are nonzero in 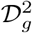. IBRE is particularly important, as our primary interest lies in determining whether these methods can accurately recover meaningful drug-CCL interactions, which we primarily characterize as “tiny blocks and stripes” in 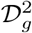.

The comparative results for three synthetic datasets are presented in Figures 1, 2, and 3. These datasets were generated using the three models for 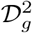 and a product of two low-rank matrices for 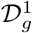 as described in the last section. Additional studies on synthetic data, utilizing a different model for 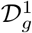 and varying block sizes, are available on GitHub.

In the case of the synthetic data with the simple two-block model for 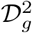 depicted in Figure 1, RT-DMF consistently surpasses all other methods in terms of recovery error (RE) and in-block recovery error (IBRE), even though the other methods also demonstrate satisfactory performance.

In the case of the synthetic data with mixed overlapping blocks for 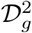, as shown in Figure 2, both MF and RPCA nearly completely fail to perform effectively, likely because the data doesn’t align with the underlying assumptions of MF and RPCA. However, this synthetic dataset represents an important real-world scenario: CCLs might respond to various drug mechanisms, leading to blocks with overlapping CCLs. Similarly, drugs that share a common mode of action could have mechanisms that overlap with other drugs, resulting in blocks with overlapping drugs. The outcomes of this dataset also imply that MSE may not be the most suitable metric for comparing methods. Despite MF and RPCA’s inability to recover any substantial information, they still manage to achieve lower test MSE compared to RT-DMF.

For the synthetic data incorporating real data residuals shown in Figure 3, RT-DMF achieves impressively low in-block recovery error, indicating its ability to recover true signals even in the presence of relatively complex noise.

These simulation studies emphasize RT-DMF’s capability to distinguish and retrieve meaningful signals from complex data across diverse scenarios. This makes RT-DMF an invaluable tool in the analysis of drug sensitivity data, where discerning subtle yet important signals is crucial for identifying potentially effective therapeutic interventions.

### 3.4 Stability Studies

Deep learning methods are often criticized for their instability (Yu and Kumbier (2020)), as they typically involve hundreds of iterations of random samples for stochastic gradient descent following a randomized initialization of weights. However, RT-DMF does not suffer from random sampling as we utilize the full gradient in each iteration. In this section, we demonstrate empirically that RT-DMF is robust with respect to initialization randomization – it can recover nearly identical matrices given different initializations with an appropriate stopping criterion.

We use the same real data residual model for 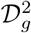 as in the previous section but vary the initialization by setting three different seeds. The results are illustrated in Figure 8 in the appendix. Although different stopping epochs are employed (choosing the epoch just before the validation MSE begins to increase), our recovered matrices show minimal differences in terms of the evaluation metrics, as demonstrated in the figure in the appendix.

## 4. REAL DATA RESULTS

We present our results from multiple perspectives to show that RT-DMF successfully retains meaningful information in real data. We apply RT-DMF to five relatively large and well-prepared datasets individually: GDSC1000, CTRPv2, FIMM, GRAY and PRISM. For each pair of the datasets, we use the overlapping part to calculate the correlation between the drug-response profiles in the two datasets for each overlapping drug. We show that the percentage of significantly correlated drugs increases after RT-DMF is applied. We then present some case studies of individual drugs on how RT-DMF can reveal meaningful information after correcting the data. We also demonstrate the validity of RT-DMF on cell-lines that have known mutations. Lastly, we demonstrate its effectiveness on a downstream analysis scenario to show that not only the correction part of RT-DMF provides meaningful information, but also the imputation part can help improve downstream tasks.

### 4.1 Increase of Significantly Correlated Drugs

We use a two-tailed Spearman’s rank correlation test presented in Zar (1972) with p-value 0.01 to determine if a particular drug’s Spearman’s rank correlation is significant across two datasets. In the table 1, we summarize the percentage of drugs that show significant Spearman’s correlation in the original and corrected datasets. For comparison, we denote the correlation before correction with a parenthesis under the correlation after correction. We observe that in general, RTDMF increases the percentage of drugs that show significant correlation across different datasets. Notably, over 20% more drugs are correlated between GDSC1000 and CTRPV2, two highly cited pharmacogenomics datasets. However, One might notice there are a few exceptions occurring at FIMM and GRAY. This is because they are very small datasets, and hence have very few over-lapping cell-lines with other datasets in general. For example, FIMM and GRAY only have 11 overlapping cell-lines for the computation of Spearman’s rank correlation. The critical value gets more stringent when the vector size gets smaller, namely two drugs need to show extremely strong Spearman correlation to be considered significantly correlated when the vector size is small. Nevertheless, for datasets that contain abundant drugs and CCL’s, such as GDSC1000, CTRPv2 and PRISM, we observe a considerable improvement trend.

**Table 1:** Percentage of drugs that show significant Spearman’s correlation before (values in parentheses) and after (values shaded) the application of RT-DMF.

|  | GDSC1000 | CTRPv2 | FIMM | GRAY | PRISM |
| --- | --- | --- | --- | --- | --- |
| GDSC1000 | - | 83.3%<br>(62.5%) | 43.8%<br>(18.8%) | 42.1%<br>(21.1%) | 47.5%<br>(35.6%) |
| CTRPv2 | - | - | 66.7%<br>(60%) | 50%<br>(35.7%) | 57.6%<br>(49.1%) |
| FIMM | - | - | - | 5.26%<br>(5.26%) | 6.67%<br>(6.67%) |
| GRAY | - | - | - | - | 28.8%<br>(18.6%) |
| PRISM | - | - | - | - | - |

### 4.2 Case Studies of Individual Drugs

We also present scatter plots for some drugs individually for a better illustration of the performance of RT-DMF in figure 4. In this set of illustrations, each data point represents the value of summarized dose-response values in the dataset labeled on axes, hence the blue line *y* = *x* represents the perfect situation that the values of such drug cell-line responses are exactly the same between two datasets. Different colors represent the value in original and RT-DMF corrected datasets. We may observe that the reason why correlation increases is because the rightmost points are well corrected by the method, as it’s much closer to the *y* = *x* line. From figure 4a, 4b and 4c, we observe that RT-DMF is able to alleviate the datapoints that form a “stripe” in the original scatterplot. Figure 4d shows an exceptional case, where most of the data points in PRISM have a number around 0 but they vary a lot in the GDSC dataset, likely due to an erroneous input in PRISM. However, we still see RTDMF attempts to adjust some values that are low in GDSC but high in PRISM.

**Fig. 4:**
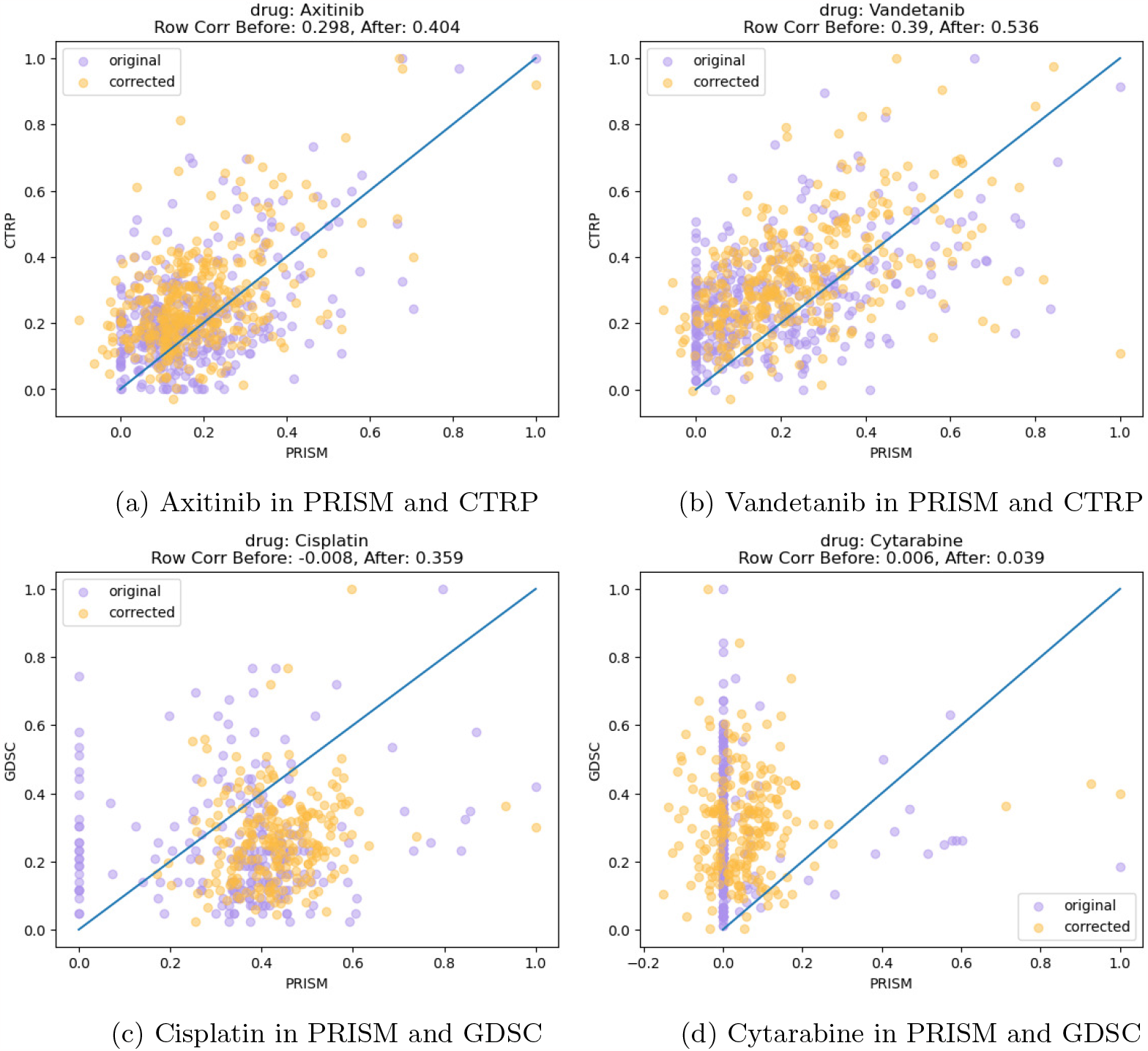
Individual drug scatter plots across different datasets. Each dot represents the value of one cell-line corresponding to such drug in the *x*-axis dataset and *y*-axis dataset.

We present a typical biomarker discovery case to demonstrate the utilization of the corrected data. It is known that BRAF mutation is a biomarker to predict the efficacy of Dabrafinib, a BRAF inhibitor; hence we expect to see a difference in Dabrafinib’s sensitivity between BRAF mutated and wild type cell lines (as denoted by 1 and 0). We observe that in figure 5, after RT- DMF is applied, the difference between the two groups in GDSC becomes even more profound compared to the original. This is specifically demonstrated that the wild type group in RT-DMF processed data is much less varied than the original data, which leads to better separation. This demonstrates that the correction of RT-DMF does provide biologically more meaningful data.

**Fig. 5:**
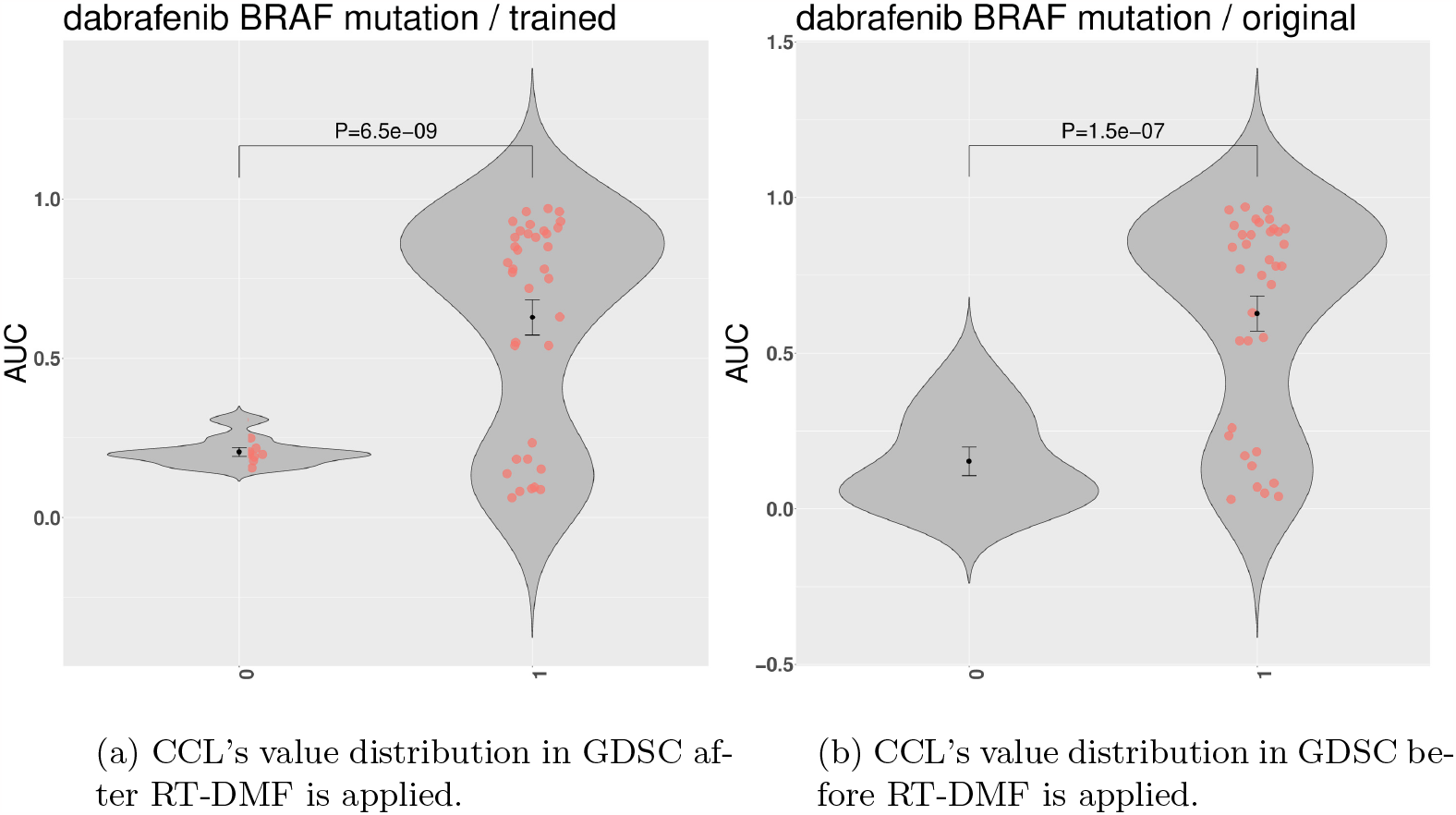
We can see that the 0 (non-mutation group) and 1 (mutation group) have a more significant difference after RT-DMF is applied to GDSC.

### 4.3 Application of Drug-Sensitivity Data on Other Tasks

*Drug Sensitivity Prediction* Gene expression profiling of cell lines is routinely performed in the lab, while testing the sensitivity of a large number of drugs in a cell line is costly and time-consuming. It would be helpful if a machine learning model could predict drug sensitivity in a cell line based on its gene expression profile. However, the model performance may be limited by the poor quality of drug sensitivity data. Our recent work surveyed commonly used machine learning in this task and found one effective method is to use an elastic net as in Yeh *and others* (2022). Since an elastic net model requires a full matrix, the missing values in the original data were imputed by KNN-Imputer before predictions as in the paper Yeh *and others* (2022). In order to find whether the data corrected and imputed by RT-DMF helps improve the prediction performance, as measured by rooted mean squared error. We apply the elastic net model to both CTRP and GDSC data and compare the prediction performance between the original data and the corrected and imputed data by RT-DMF. We find that the data processed by RT-DMF decreases the rooted mean squared error from 0.023 to 0.009 in CTRP, 0.047 to 0.031 in GDSC and also reduces the variation as in Figure 6, suggesting that the corrected data could better serve the downstream tasks.

**Fig. 6:**
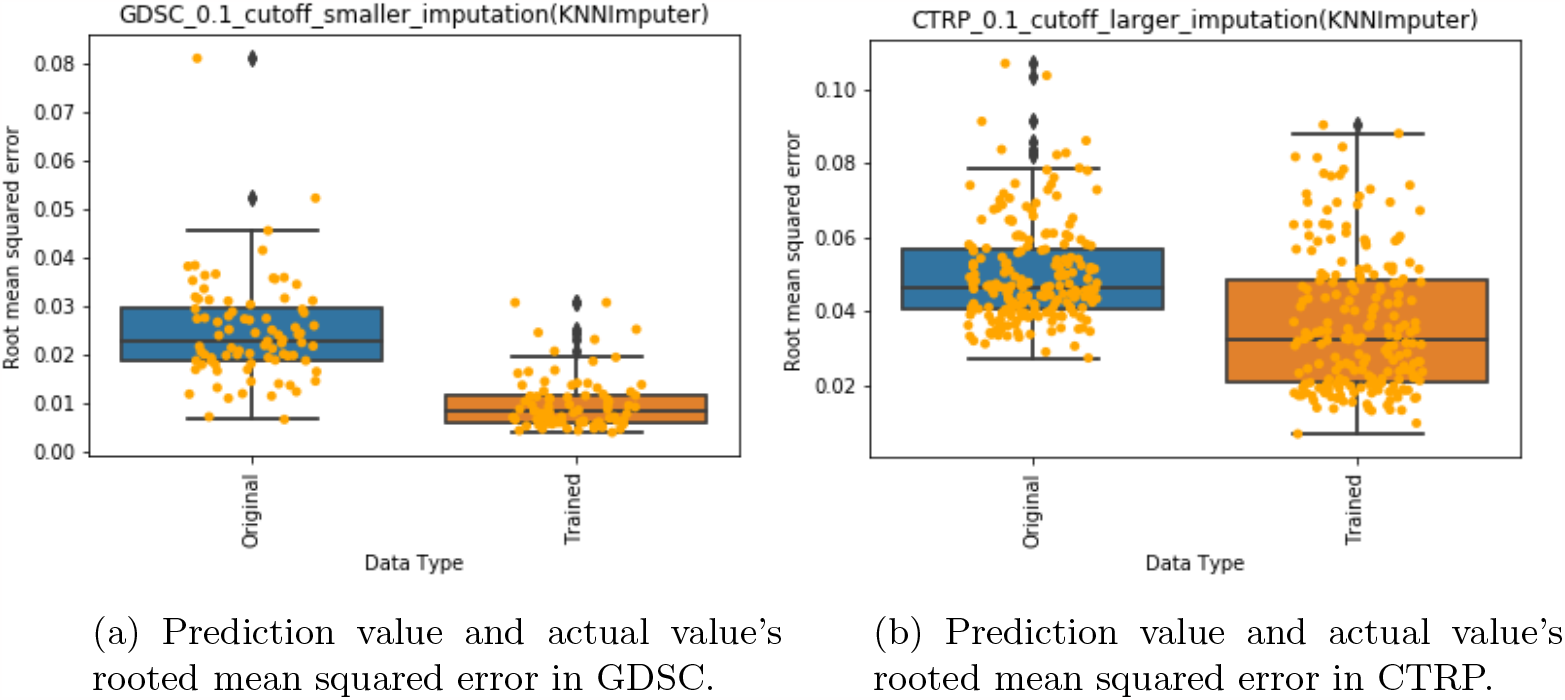
RT-DMF trained datasets provide better results in terms of prediction in terms of RMS.

*Clustering Analysis* Another application of drug sensitivity data lies in providing insights into the relations among drugs or cell lines via clustering analysis (Wang and others). Such an analysis facilitates the grouping of drugs sharing similar Mechanisms of Action (MOA), which can foster the identification of alternative treatments, as well as facilitate the discovery of new drugs and targets. On the other hand, clustering cell lines with similar drug response patterns creates opportunities for exploring underlying biological mechanisms and discovering potential personalized therapies by predicting individual patient responses using cell line data.

To demonstrate that RT-DMF could improve clustering accuracy, we performed clustering analysis using RT-DMF corrected and the original drug sensitivity data from GDSC and CTRP. The missing values in the original data were imputed by KNN-Imputer before clustering as in Yeh *and others* (2022). We compared the clusters of drugs / cell lines to their pre-defined categories, and quantitatively evaluate the clustering performance by computing cluster accuracy (CA) as in Fahad *and others* (2014).

For the drug clustering, we utilized GDSC drugs that had pre-defined MOAs associated with EGFR signaling and Metabolism, and CTRP drugs whose MOAs were defined as inhibitors of IKK-2 and gamma-secretase. These paths and proteins are distinctly characterized by different functions, mechanisms, and associated diseases. EGFR signaling primarily regulates cell growth, proliferation, and survival through signal transduction cascades, whereas Metabolism plays a role in energy production, molecule biosynthesis, and cellular homeostasis. Similarly, while IKK- 2 primarily controls NF-*κ*B signaling and participates in inflammation and immune responses, gamma-secretase is responsible for proteolytic processing of transmembrane proteins, playing a significant role in A*β* production (associated with Alzheimer’s disease) and Notch signaling. Given this distinctiveness, we anticipated that these selected drugs would display unique patterns between categories. Our findings confirmed this, as we observed a CA of 0.571 for the original GDSC drug data and 0.714 for the RT-DMF corrected data, and a CA of 1 for both original and corrected CTRP drug data as demonstrated in 7A and 7B.

For cell line clustering, we used GDSC CCLs sourced from chondrosarcoma, chronic myeloid leukemia, and B cell leukemia, and CTRP CCLs sourced from acute lymphoblastic B cell leukemia and Grade IV astrocytoma. These represent distinctly different cancers with diverse cell and tissue origins. Acute lymphoblastic B cell leukemia is a blood cancer originating from B cells, whereas Grade IV astrocytoma is a brain tumor originating from astrocytes. Similarly, chondrosarcoma is a bone cancer that begins in chondrocytes, chronic myeloid leukemia arises from myeloid cells, and B cell leukemia originates from B cells. Based on these differences, we anticipated unique patterns between these CCLs. This expectation was met with a CA of 1 for both original and RT-DMF corrected CTRP data, and a CA of 0.630 for original GDSC data and 0.704 for the RT-DMF corrected data as demonstrated in figure 7C and 7D.

**Fig. 7:**
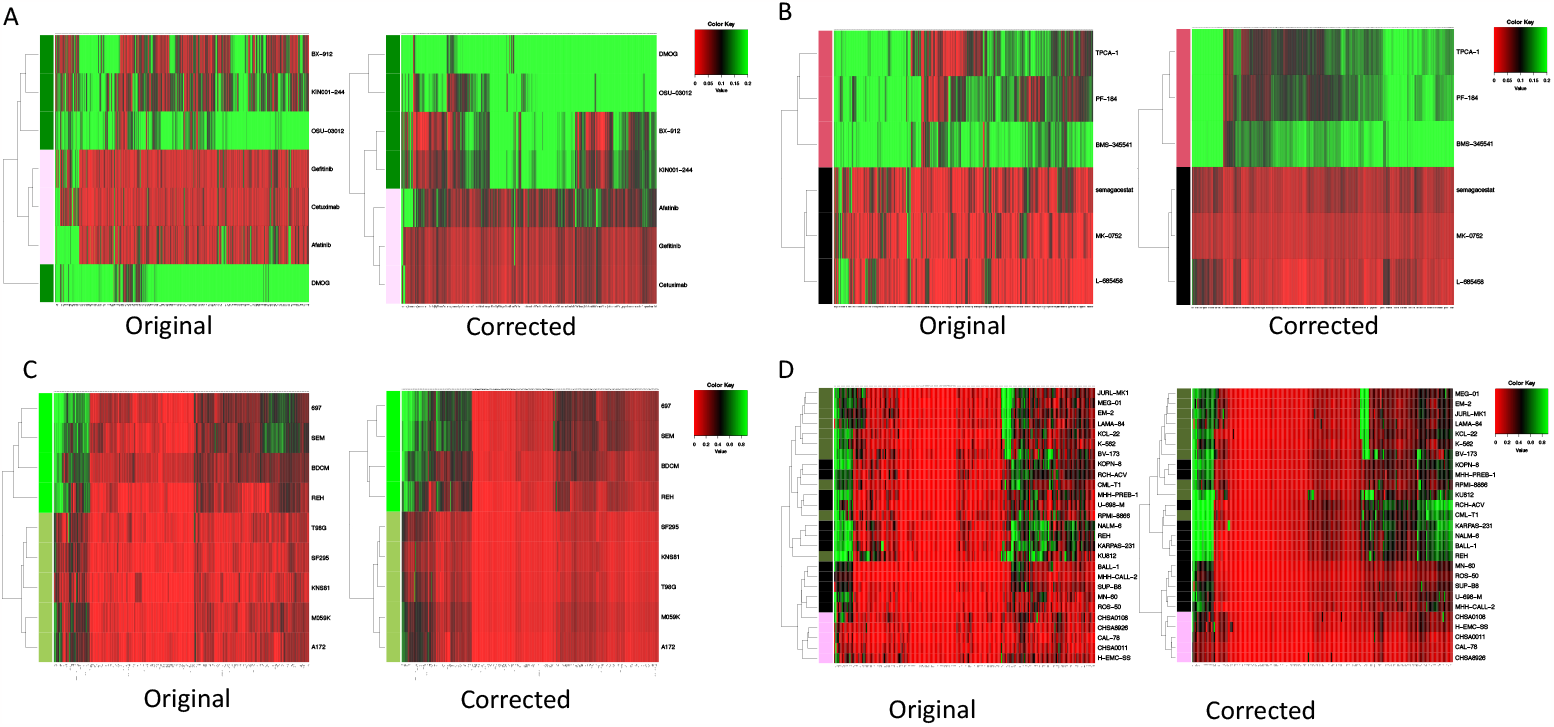
Heatmaps colored by the drug sensitivity values show the clustering results. Left: Using the original data. Right: Using the RT-DMF corrected data. a, clustering of selected GDSC drugs. b, clustering of selected CTRP drugs. c, clustering of selected CTRP cell lines. d, clustering of selected GDSC cell lines.

**Fig. 8:**
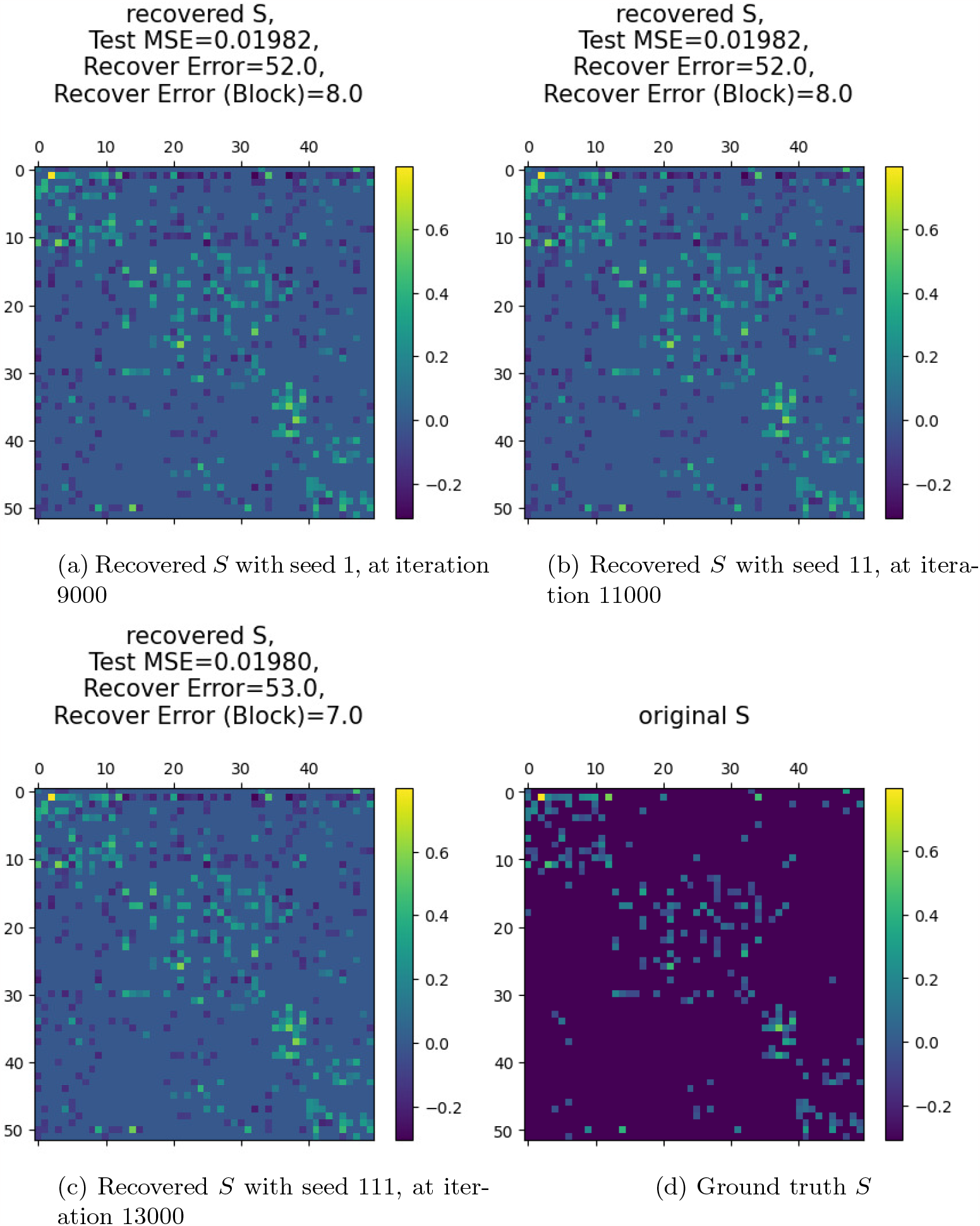
Replicate of recovering *S* using different initialization with the same stop criterion – the last epoch where validation MSE starts to rise.

The analysis shows an enhancement in clustering accuracy when the RT-DMF corrected drug sensitivity data is applied, compared to the original data. This implies that data correction via the RT-DMF method holds significant potential for improving downstream tasks, thereby enriching the impact and outcomes

## FUNDING

This work is supported by the National Institute Of General Medical Sciences of the National Institutes of Health under Award Number R01GM134307. The content is solely the responsibility of the authors and does not necessarily represent the official views of the National Institutes of Health.

## APPENDIX Implementation Details

We discuss about some implementation details of RT-DMF and some other methods we used to bench-mark, specifically, we discuss the meaning of parameters and why a good 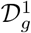 will lead to a good 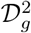. In the sections we also discuss the data synthesis details.

One might notice that the neural network in RT-DMF is a wide one. In fact, the width does not violate the low-rank assumption. This is due to an implicit regularization during the optimization of this model. What’s more, such a linear DMF is scalable and surprisingly stable to initialization as long as the stopping criterion is consistent.

In fact, the tuning parameters in all three methods can be seen as a trade-off between the fitting of an appropriate low-rank 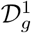: we want 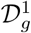 to be informatically similar to the original matrix in terms of intrinsic properties of drugs and CCLs and their interactions. A poor fitting of 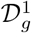 will lead to the hardness in determining appropriate 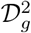, hence losing the potential therapeutically information that biologists are seeking for. As what we have observed in the 3, other matrix completion methods are not successful in terms of recovering 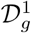 and 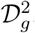.

The early stopping step helps the determination of the 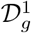 and 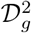 under the consideration / assumption that general interactions between drugs and cancer cell lines or systematic structure / pattern, which are expected to present in 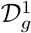, are less likely interesting therapeutically. In contrast, 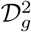 consists of information reflecting more specific interactions between drugs and cancer cell lines which are more likely to be therapeutically relevant.

The thresholding step helps better separate 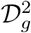 and Σ using magnitude differences and enrichment in pre-known drug and CCL groups.

After recovering, we do not know which is the signal and which is the noise, that is: it’s hard to discriminate between 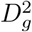 and Σ. However, though 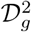 would be more of interest, a poor recovery of 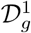 will lead to a poor discovery of 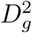. As specifically, we only consider the drug-cell interaction data and there is no other information when training the model. Hence our 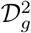 can be only recovered by the residual information between the original dataset and 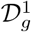. The low ranked structure corresponding to larger (more general effect) will be less likely to be interesting and the smaller (more specific group design) will be more likely to be interesting.

*Generation Details of* 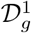 We generate 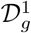 in two different ways:

- As a product of two nonnegative low-rank matrices: we first generate a matrix *L* where each component *L*_*ij*_ is uniformly sampled from [0, 1], next we perform nonnegative matrix factorization on *L* as in Pedregosa *and others* (2011), i.e., *L ≈ U*_NMF_*V*N^*?*^MF, where *U*_NMF_, *V*_NMF_ are low-rank matrix with rank 10. Then we let 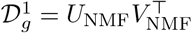.
- As a product of multiple non-negative low rank matrices: we first generate a matrix *L* where each component *L*_*ij*_ is uniformly sampled from [0, 1], next we perform deep matrix factorization on *L* with four matrix components, i.e., *L ≈ W*_1_*W*_2_*W*_3_*W*_4_, where *W*_1_, *W*_2_, *W*_3_, *W*_4_ are low-rank matrices with rank found by DMF. Then we let 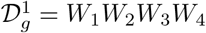.
- We used the 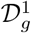 generated by the first method in the simulation experiments, as we do not observe much differences between the two.

*Generation Details of* 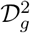 *under Simple Two Block Assumptions* First we generate an all-zero ma-

trix. Then, for entries 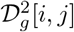 such that *i, j ∈* [*{r*_1_, …, *r*_bsize_*}×{c*_1_, *c*_2_, …, *c*_bsize_*}*]*∪*[*{r*_bsize+1_, …, *r*_2*·*bsize_*}× {c*_bsize+1_, …, *c*_2*·*bsize_*}*] (in block, and bsize denotes the block size), we have *D*_*l*_[*i, j*] have a probability of *p*_inside_ being magnitude of *s ∈* Unif(0.2, 0.3). We pick such an interval because it is slightly larger than 2 standard deviations above average for 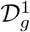. Out-block entries have a probability of *p*_outside_ = 0.05 being magnitude of *s* = 0.3 to replace the 0 value. We set *p*_inside_ = 0.6, *p*_outside_ = 0.05, *s* = 0.3 across all block sizes.

*Generation Details of* 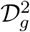 *under Mixed Overlapping Block Assumptions* We generate an all-zero matrix first again. Then for entries *D*^2^[*i, j*] such that *i, j ∈ {*[*{r*_1_, …, *r*_20_*} × {c*_1_, *c*_2_, …, *c*_10_*}*] *∪* [*{r*_5_, *r*_6_, …, *r*_20_*} × {c*_25_, …, *c*_35_*}*] *∪* [*{r*_30_, …, *r*_35_*} × {c*_0_, *c*_1_, …, *c*_10_*}*] *∪* [*{r*_0_, …, *r*_40_*} × c*_40_*}*] (in block), we have 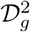 have a probability of *p*_inside_ being magnitude of *s*, while out-block entries have a probability of *p*_outside_ being magnitude of *s* to replace the original 0. We set *p*_inside_ = 0.6, *p*_outside_ = 0.05, *s* = 0.3

*Generation Details of* 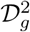 *under Real Data Assumptions* This model directly takes the residual from the FIMM data in Mpindi *and others* (2016) and its naive low-rank representation. In particular, we find a naive low-rank representation of FIMM data and fetch the residual through subtracting the real data from the low-rank data entry-wise. We then do a simple bi-clustering to the residual through spectral methods. We select the found bi-clusters that are most significant in terms of in-group size and take them as true signals. The rest of the entries have a certain probability *p* to be kept. All other entries are set to 0 so that we assume they are irrelevant noises. This is a simplified version of real data noise as we assume the residual that matters only lies in the groups we have found. An visualization of *D*^2^ under this situation when *p* = 0.1 is demonstrated in figure 3d.

## Notes

https://github.com/tomwhoooo/rtdmf

### Competing Interest Statement

The authors have declared no competing interest.

## References

Arora, Sanjeev, Cohen, Nadav, Hu, Wei and Luo, Yuping. (2019). Implicit Regularization in Deep Matrix Factorization. Red Hook, NY, USA: Curran Associates Inc.

Barretina, J., Caponigro, G., Stransky, N., Venkatesan, K., Margolin, A. A., Kim, S., Wilson, C. J., Lehar, J., Kryukov, G. V., Sonkin, D., Reddy, A., Liu, M., Murray, L., Berger, M. F., Monahan, J. E., Morais, P., Meltzer, J., Korejwa, A., Jane-Valbuena, J., Mapa, F. A., Thibault, J., Bric-Furlong, E., Raman, P., Shipway, A., Engels, I. H., Cheng, J., Yu, G. K., Yu, J., Aspesi, P., de Silva, M., Jagtap, K., Jones, M. D., Wang, L., Hatton, C., Palescandolo, E., Gupta, S., Mahan, S., Sougnez, C., Onofrio, R. C., Liefeld, T., MacConaill, L., Winckler, W., Reich, M., Li, N., Mesirov, J. P., Gabriel, S. B., Getz, G., Ardlie, K., Chan, V., Myer, V. E., Weber, B. L., Porter, J., Warmuth, M., Finan, P., Harris, J. L., Meyerson, M., Golub, T. R., Morrissey, M. P., Sellers, W. R., Schlegel, R. and others. (2012, Mar). The Cancer Cell Line Encyclopedia enables predictive modelling of anticancer drug sensitivity. Nature 483(7391), 603–607.

Basu, A., Bodycombe, N. E., Cheah, J. H., Price, E. V., Liu, K., Schaefer, G. I., Ebright, R. Y., Stewart, M. L., Ito, D., Wang, S., Bracha, A. L., Liefeld, T., Wawer, M., Gilbert, J. C., Wilson, A. J., Stransky, N., Kryukov, G. V., Dancik, V., Barretina, J., Garraway, L. A., Hon, C. S., Munoz, B., Bittker, J. A., Stockwell, B. R., Khabele, D., Stern, A. M., Clemons, P. A., Shamji, A. F. and others. (2013, Aug). An interactive resource to identify cancer genetic and lineage dependencies targeted by small molecules. Cell 154(5), 1151–1161.

Bouhaddou, Mehdi, DiStefano Matthew S., Riesel Eric A., Carrasco, Emilce, Holzapfel Hadassa Y., Jones DeAnalisa C., Smith Gregory R., Stern, Alan D., Somani Sulaiman S., Thompson, T. Victoria and others. (2016, 11). Drug response consistency in ccle and cgp. Nature 540, E9 EP –.

Candès Emmanuel J., Li, Xiaodong, Ma, Yi and Wright, John. (2011, jun). Robust principal component analysis? J. ACM 58(3).

Candès Emmanuel J and Recht, Benjamin. (2009). Exact matrix completion via convex optimization. Foundations of Computational mathematics 9(6), 717–772.

Chen, B. and Butte, A. J. (2016, Mar). Leveraging big data to transform target selection and drug discovery. Clin. Pharmacol. Ther. 99(3), 285–297.

Collins, F. S. and Varmus, H. (2015, Feb). A new initiative on precision medicine. N. Engl. J. Med. 372(9), 793–795.

Daemen, A., Griffith, O. L., Heiser, L. M., Wang, N. J., Enache, O. M., Sanborn, Z., Pepin, F., Durinck, S., Korkola, J. E., Griffith, M., Hur, J. S., Huh, N., Chung, J., Cope, L., Fackler, M. J., Umbricht, C., Sukumar, S., Seth, P., Sukhatme, V. P., Jakkula, L. R., Lu, Y., Mills, G. B., Cho, R. J., Collisson, E. A., van’tVeer, L. J., Spellman, P. T. and others. (2013). Modeling precision treatment of breast cancer. Genome Biol. 14(10), R110.

de Gramont, A., Watson, S., Ellis, L. M., Rodon, J., Tabernero, J., de Gramont, A. and Hamilton, S. R. (2015, Apr). Pragmatic issues in biomarker evaluation for targeted therapies in cancer. Nat Rev Clin Oncol 12(4), 197–212.

Fahad, Adil, Alshatri, Najlaa, Tari, Zahir, Alamri, Abdullah, Khalil, Ibrahim, Zomaya Albert Y., Foufou, Sebti and Bouras, Abdelaziz. (2014). A survey of clustering algorithms for big data: Taxonomy and empirical analysis. IEEE Transactions on Emerging Topics in Computing 2(3), 267–279.

Garnett, M. J. and et al. (2012, Mar). Systematic identification of genomic markers of drug sensitivity in cancer cells. Nature 483(7391), 570–575.

Hafner, M., Niepel, M., Chung, M. and Sorger, P. K. (2016, 06). Growth rate inhibition metrics correct for confounders in measuring sensitivity to cancer drugs. Nat. Methods 13(6), 521–527.

Haibe-Kains, B., El-Hachem, N., Birkbak, N. J., Jin, A. C., Beck, A. H., Aerts, H. J. and Quackenbush, J. (2013, Dec). Inconsistency in large pharmacogenomic studies. Nature 504(7480), 389–393.

Hu, Zhiyue Tom, Ye, Yuting, Newbury Patrick A., Huang, Haiyan and Chen, Bin. (2019). AICM: A Genuine Framework for Correcting Inconsistency Between Large Pharmacogenomics Datasets . pp. 248–259.

Iorio, F., Knijnenburg, T. A., Vis, D. J., Bignell, G. R., Menden, M. P., Schubert, M., Aben, N., Goncalves, E., Barthorpe, S., Lightfoot, H., Cokelaer, T., Greninger, P., van Dyk, E., Chang, H., de Silva, H., Heyn, H., Deng, X., Egan, R. K., Liu, Q., Mironenko, T., Mitropoulos, X., Richardson, L., Wang, J., Zhang, T., Moran, S., Sayols, S., Soleimani, M., Tamborero, D., Lopez-Bigas, N., Ross-Macdonald, P., Esteller, M., Gray, N. S., Haber, D. A., Stratton, M. R., Benes, C. H., Wessels, L. F. A., Saez-Rodriguez, J., McDermott, U. and others. (2016, Jul). A Landscape of Pharmacogenomic Interactions in Cancer. Cell 166(3), 740–754.

Lee, S. I., Celik, S., Logsdon, B. A., Lundberg, S. M., Martins, T. J., Oehler, V. G., Estey, E. H., Miller, C. P., Chien, S., Dai, J., Saxena, A., Blau, C. A. and others. (2018, 01). A machine learning approach to integrate big data for precision medicine in acute myeloid leukemia. Nat Commun 9(1), 42.

Lowy, D. R. and Collins, F. S. (2016, May). Aiming High–Changing the Trajectory for Cancer. N. Engl. J. Med. 374(20), 1901–1904.

Marcotte, R., Sayad, A., Brown, K. R., Sanchez-Garcia, F., Reimand, J., Haider, M., Virtanen, C., Bradner, J. E., Bader, G. D., Mills, G. B., Pe’er, D., Moffat, J. and others. (2016, Jan). Functional Genomic Landscape of Human Breast Cancer Drivers, Vulnerabilities, and Resistance. Cell 164(1-2), 293–309.

Mpindi, John Patrick, Yadav, Bhagwan, Östling Päivi, Gautam, Prson, Malani, Disha Murumägi, Astrid, Hirasawa, Akira, Kangaspeska, Sara, Wennerberg, Krister, Kallioniemi, Olli and others. (2016, 11). Consistency in drug response profiling. Nature 540, E5 EP –.

Neyshabur, Behnam. (2017). Implicit regularization in deep learning. CoRR abs/1709.01953.

Niepel, M., Hafner, M., Pace, E. A., Chung, M., Chai, D. H., Zhou, L., Schoeberl, B. and Sorger, P. K. (2013, Sep). Profiles of Basal and stimulated receptor signaling networks predict drug response in breast cancer lines. Sci Signal 6(294), ra84.

Pedregosa, F., Varoquaux, G., Gramfort, A., Michel, V., Thirion, B., Grisel, O., Blondel, M., Prettenhofer, P., Weiss, R., Dubourg, V., Vanderplas, J., Passos, A., Cournapeau, D., Brucher, M., Perrot, M. and others. (2011). Scikit-learn: Machine learning in Python. Journal of Machine Learning Research 12, 2825–2830.

Rees, M. G., Seashore-Ludlow, B., Cheah, J. H., Adams, D. J., Price, E. V., Gill, S., Javaid, S., Coletti, M. E., Jones, V. L., Bodycombe, N. E., Soule, C. K., Alexander, B., Li, A., Montgomery, P., Kotz, J. D., Hon, C. S., Munoz, B., Liefeld, T., Dan?ik, V., Haber, D. A., Clish, C. B., Bittker, J. A., Palmer, M., Wagner, B. K., Clemons, P. A., Shamji, A. F. and others. (2016, Feb). Correlating chemical sensitivity and basal gene expression reveals mechanism of action. Nat. Chem. Biol. 12(2), 109–116.

Safikhani, Z., Smirnov, P., Freeman, M., El-Hachem, N., She, A., Rene, Q., Goldenberg, A., Birkbak, N. J., Hatzis, C., Shi, L., Beck, A. H., Aerts, H. J. W. L., Quackenbush, J. and others. (2016). Revisiting inconsistency in large pharmacogenomic studies. F1000Res 5, 2333.

Seashore-Ludlow, B., Rees, M. G., Cheah, J. H., Cokol, M., Price, E. V., Coletti, M. E., Jones, V., Bodycombe, N. E., Soule, C. K., Gould, J., Alexander, B., Li, A., Montgomery, P., Wawer, M. J., Kuru, N., Kotz, J. D., Hon, C. S., Munoz, B., Liefeld, T., Dan?ik, V., Bittker, J. A., Palmer, M., Bradner, J. E., Shamji, A. F., Clemons, P. A. and others. (2015, Nov). Harnessing Connectivity in a Large-Scale Small-Molecule Sensitivity Dataset. Cancer Discov 5(11), 1210–1223.

Wang, S., Huang, E., Cairns, J., Peng, J., Wang, L. and Sinha, S. Identification of pathways associated with chemosensitivity through network embedding. PLoS computational biology 15(3).

Yang, W., Soares, J., Greninger, P., Edelman, E. J., Lightfoot, H., Forbes, S., Bindal, N., Beare, D., Smith, J. A., Thompson, I. R., Ramaswamy, S., Futreal, P. A., Haber, D. A., Stratton, M. R., Benes, C., McDermott, U. and others. (2013, Jan). Genomics of Drug Sensitivity in Cancer (GDSC): a resource for therapeutic biomarker discovery in cancer cells. Nucleic Acids Res. 41(Database issue), D955–961.

Yeh, Shan-Ju, Chen, Ruoqiao, Xing, Jing, Sun, Mengying, Liu, Ke, Paithankar, Shreya, Zhou, Jiayu and Chen, Bin. (2022, 06). Transcell: In silico characterization of genomic landscape and cellular responses from gene expressions through a two-step transfer learning. Cancer Research 82(12), 1927–1927.

Yothers, G., O’Connell, M. J., Lee, M., Lopatin, M., Clark-Langone, K. M., Mill-ward, C., Paik, S., Sharif, S., Shak, S. and Wolmark, N. (2013, Dec). Validation of the 12-gene colon cancer recurrence score in NSABP C-07 as a predictor of recurrence in patients with stage II and III colon cancer treated with fluorouracil and leucovorin (FU/LV) and FU/LV plus oxaliplatin. J. Clin. Oncol. 31(36), 4512–4519.

Yu, Bin and Kumbier, Karl. (2020). Veridical data science. Proceedings of the National Academy of Sciences 117(8), 3920–3929.

Zar, Jerrold H. (1972). Significance testing of the spearman rank correlation coefficient. Journal of the American Statistical Association 67, 578–580.

